# A Fluorescence-based High Throughput-Screening assay for the SARS-CoV RNA synthesis complex

**DOI:** 10.1101/2020.07.07.192005

**Authors:** Cecilia Eydoux, Veronique Fattorini, Ashleigh Shannon, Thi-Tuyet-Nhung Le, Bruno Didier, Bruno Canard, Jean-Claude Guillemot

## Abstract

The Severe Acute Respiratory Syndrome Coronavirus (SARS-CoV) emergence in 2003 introduced the first serious human coronavirus pathogen to an unprepared world. To control emerging viruses, existing successful anti(retro)viral therapies can inspire antiviral strategies, as conserved viral enzymes (eg., viral proteases and RNA-dependent RNA polymerases) represent targets of choice. Since 2003, much effort has been expended in the characterization of the SARS-CoV replication/transcription machinery. Until recently, a pure and highly active preparation of SARS-CoV recombinant RNA synthesis machinery was not available, impeding target-based high throughput screening of drug candidates against this viral family. The current Severe Acute Respiratory Syndrome Coronavirus-2 (SARS-CoV-2) pandemic revealed a new pathogen whose RNA synthesis machinery is highly (>96% aa identity) homologous to SARS-CoV. This phylogenetic relatedness highlights the potential use of conserved replication enzymes to discover inhibitors against this significant pathogen, which in turn, contributes to scientific preparedness against emerging viruses. Here, we report the use of a purified and highly active SARS-CoV replication/transcription complex (RTC) to set-up a high-throughput screening of Coronavirus RNA synthesis inhibitors. The screening of a small (1,520 compounds) chemical library of FDA-approved drugs demonstrates the robustness of our assay and will allow to speed-up drug repositioning or novel drug discovery against the SARS-CoV-2.

**Principle of SARS-CoV RNA synthesis detection by a fluorescence-based high throughput screening assay:** 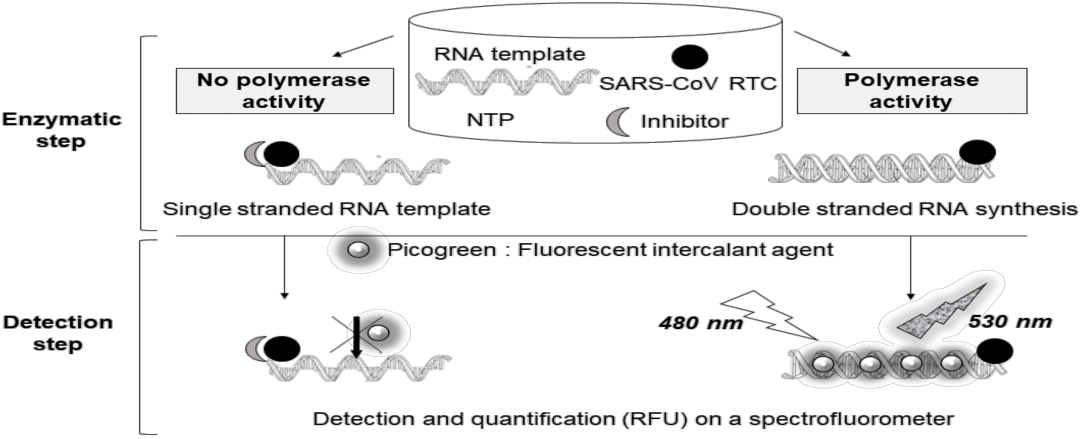

**Highlights:**

- A new SARS-CoV non radioactive RNA polymerase assay is described
- The robotized assay is suitable to identify RdRp inhibitors based on HTS

## 1. Introduction

The current worldwide Corona Virus Disease 2019 (COVID-19) crisis highlights the urgent need of cost-effective drugs able to control the SARS-CoV-2 at the earliest stage of infection (Wu et al., 2020; Zhou et al., 2020). Such a drug would find its usefulness immediately after a patient’s positive SARS-CoV-2 diagnostics, to help quarantine effectiveness and acceptance, control clusters of cases, reduce potentially severe outcomes, and ultimately curb the epidemics. A huge effort is being pursued across the world to find drugs against the SARS-CoV-2 virus. Towards this endeavor, one of the most potentially rapid ways is to screen chemical libraries of FDA-approved compounds. Under favorable conditions, an approved drug can be compatible with its novel antiviral indication, and thus go under accelerated clinical evaluation (Cheng et al., 2019).

Antiviral drug discovery can be conducted through several screening methods, each having its advantages and drawbacks. Phenotypic screening makes use of virus infected cells which are rescued when a compound prevents or dampens the consequences of the infection, as judged by cytopathic effect or virus production read-outs. In this type of process, the target of the identified hit is not known, and thus, hit expansion is conducted in a somewhat blind manner. Several methods exist to determine the viral or cellular target, allowing potential rapid hit-to-lead development implemented in conjunction with structural biology techniques and functional assays. Amongst the advantages of this method, drug uptake by the cell, metabolic activation/decay, and primary bioavailability and toxicity features are included in the screening process.

Target-based screening generally proposes to select ligands or inhibitors through the use of the purified target known to play a critical role for pathogen propagation. Besides the insurance of knowing the actual inhibitor target, the immediate advantage is that screening can be performed in standard BSL-1 or 2 laboratories, and hit expansion is immediately and greatly facilitated by functional assays, often guided by available structural data on the target of interest. Obvious disadvantages are that an excellent inhibitor may never cross cellular membranes nor reach its target in the cell.

Clearly, both types of screening methods are required, and even more importantly, they are complementary to expedite drug design, from discovery to hit-to-lead optimization.

Viral RNA-dependent RNA polymerases (RdRp) represent an extremely attractive target for antiviral drugs due to their high level of sequence and structural conservation and vital role in viral replication (Venkataraman et al., 2018). Furthermore, they do not have any cellular homologues in the eukaryotic cell, which means that virus specific inhibition and drug selectivity can be achieved. This has been beautifully demonstrated by, amongst others, the design of Sofosbuvir, a nucleotide analogue occupying the foremost position in the fight against the hepatitis C virus (HCV). The later virus has been the subject of intense antiviral research due to its prevalence worldwide, and an expected flourishing market for antiviral drugs concomitant to various difficulties around vaccine design (Ji et al., 2019).

The situation is blatantly different for SARS-CoV, whose drug market has remained difficult to evaluate after its emergence in 2002 and subsequent vanishing in the middle of year 2003 (Pruijssers and Denison, 2019). Another reason for the lagging research of SARS-CoV is its sophisticated and multi-subunit RNA synthesis machinery (Snijder et al., 2003). It was only in 2014 that nsp7 and nsp8 were identified as co-factors of the RdRp core nsp12 and shown to confer full activity and processivity to nsp12 (Subissi et al., 2014).

These finding were corroborated by the Cryo-EM structure determination of the nsp7-8-12 SARS-CoV RNA synthesis complex in 2019, followed by that of SARS-CoV-2 in 2020 (Gao et al., 2020; Kirchdoerfer and Ward, 2019). The structure shows the nsp12 RdRp core bound to one molecule of nsp7 and two molecules of nsp8. Furthermore, the structure described by Hillen et al. suggest that nsp8 acts as a “sliding pole” for processivity of RNA synthesis (Hillen et al., 2020). Although it is not known if this represent one of the biologically relevant form(s) of the SARS-CoV Replication Transcription Complex (RTC), it does suggest that there is both biochemical and functional relevance to consider the nsp7-8-12 complex as the minimum RTC required for synthesis of the coronavirus ∼30,000 nt RNA genome.

The SARS-CoV and SARS-CoV-2 viruses are highly homologous, with a 79% overall amino acid sequence identity along the genome (Wang et al., 2020). Remarkably, the SARS-CoV nsp12 amino acid sequence is 96 % identical to that of SARS-CoV-2, and amino acid polymorphisms involve conservative changes. The genome is translated into two large polyproteins originating from Orf1a and Orf1b, the latter being expressed from a ribosomal frameshift occurring at the end of Orf1a. Orf1b codes for a set of five conserved replication proteins, among which nsp12 carries the polymerase activity when supplemented with Orf1a products nsp7 and nsp8. In the core polymerase domain, amino acid differences between SARS-CoV-1 and 2 map to the surface of the protein, not onto any of the conserved motif A to G, indicating that SARS-CoV and SARS-CoV-2 RNA synthesis properties should be nearly identical (Shannon et al., 2020).

In this paper, we use the nsp7, nsp8, and nsp12 proteins of the SARS-CoV to assemble a highly active RNA synthesis complex. We optimize reaction conditions to set-up a non-radioactive nucleotide polymerization assay with a signal-to-noise ratio appropriate for a robotized high-throughput screening (HTS) assay. We validate this HTS assay through a screening campaign using the Prestwick Chemical Library® of 1,520 FDA-EMA approved compounds, and report detailed inhibition profiles of two series of hits.

## 2. Materials and methods

### 2.1 Products and reagents

Homopolymeric adenine (Poly (A)) RNA template was obtained from GE Healthcare. Hinokiflavone (ref 1017), Apigenine (ref 1102S) and Amentoflavone (ref 1057S) were purchased from Extrasynthese and from SigmaAldrich for Quercetin (ref Q4951). Compounds were resuspended in 100% DMSO at 20 mM and stored à −20°C. 3’dUTP (ref N-3005) was purchased from Trilink. Quant-it Picogreen® dsDNA assay kit (ref P11496) was obtained from Thermofisher scientific.

### 2.2 Cloning, expression and proteins purification

The fusion protein nsp7-nsp8 was generated by inserting a GSGSGS linker sequence between the nsp7 and nsp8 coding sequences, and is named nsp7L8 (Subissi et al., 2014). The nsp7L8 and nsp12 proteins were produced and purified independently as described previously in a bacterial expression system free T7-bacteriophage RNA polymerase. The complex was reconstituted with a 1:10 ratio of nsp12:nsp7L8 as indicated.

### 2.3 Setting up of the RTC activity experimental conditions based on a fluorescent readout

#### 2.3.1 Determination of the Poly(A) template concentration

Picogreen kinetic assay was based on polymerase activity of SARS nsp12 in complex with nsp7L8, which catalyzed the reaction using a poly (A) template and uridine triphosphate (UTP). The reaction (160 µl) was carried out at 30^°^C in 50 mM Hepes pH 8, 10 mM DTT, 10 mM KCl, 2 mM MgCl_2_, 2 mM MnCl_2_, 500 µM UTP and 150 nM nsp12 as final concentrations. The final Poly(A) concentration varied from 5 nM to 500 nM. To reconstitute an active replicase, nsp7L8 was used in a 10-fold molar concentration excess compared to nsp12 as described. Nsp12 and nsp7L8 were incubated for 5 min in the presence of Poly(A) before starting the reaction with UTP addition.

30 µl of Ethylenediaminetetraacetic acid (EDTA) 100 mM were added into each well of a 96 wells black flat bottom plate (Greiner Bio-One ref 675076). At each time interval (5;7.5;10;15;20;30 and 45 min), 20 µl of reaction was added into wells on the plate with EDTA to stop the reaction. The plate was then incubated 5 minutes in the dark with 1/800 Picogreen® in TE buffer (10 mM Tris-HCl, 1 mM EDTA, pH 7.5). The plate was read on Tecan Safire II using software Magellan 6 (excited light at 480 nm, emitted light at 530 nm, optimal gain).

The assay was done in triplicate. Velocity values of each condition were determined using the Prism software by calculating the slope of the linear phase and then plotted against the Poly(A) template concentration to determine the Km (Poly(A)) and V_max_ values by using Michaelis-Menten fitting.

#### 2.3.2 Optimizing UTP concentration

UTP variation assay was performed in the same conditions as described above, with the defined Poly(A) optimum concentration of 350 nM, except for the final UTP concentration, which varied from 50 µM to 1 mM. At each time interval (5;7.5;10;15;20; 30 and 45 min), 20 µl of reaction was added into wells on the plate with EDTA to stop the reaction. The assay was experimented three times., Velocity values of each condition were determined using the Prism software by calculating the slope of the linear phase and then plotted against the UTP concentration to determine the apparent Km app (UTP) value by using Michaelis-Menten fitting or Hill fitting.

#### 2.3.3 Optimizing enzyme concentration

Enzyme variation assays were performed in the same conditions as described above (Poly(A) 350 nM, UTP 500 µM) except for the final enzyme concentration, which varied from 20 nM to 250 nM. Background and optimal gain values (given by TecanSafire 2) of each assay were determined and analyzed, using Prism software, to obtain the final best condition.

#### 2.3.4 Time course of the SARS-COV RTC assay

To assess the optimized conditions of the polymerase activity of the SARS nsp12-nsp7L8 complex, a reaction time course was performed using 350 nM poly (A) template, 500 µM UTP and 150 nM nsp12 enzyme, testing nsp12 alone, nsp7L8 alone or the nsp12/nsp7L8 complex. Nsp7L8 is in a 10-fold molar concentration excess compared to nsp12 concentration. Nsp12 and nsp7L8 were incubated at room temperature for 5 min in the presence of Poly (A) before starting the reaction by the addition of UTP. As a comparison, the assay was performed with the DV-2 polymerase (100 nM Poly (U) template, 300 µM ATP, 10 nM Enzyme as already described (Benmansour et al., 2016). Experiments, performed in triplicate, were analyzed using the Prism 5 software.

### 2.4 Robotization of the RTC assay

The assay was performed in 96-well Nunc plates. The chemical library is from Prestwick Chemical. The 1520 compounds are distributed in 19 plates, with 80 compounds per plate and the first and 12^th^ columns with DMSO. Each of the 80 compounds were added to the reaction mix using a BioMek I5 workstation (Beckman) to a final concentration of 20 µM in 5% DMSO. Reactions were conducted in 40 μL final volume. The enzyme mix containing both nsp12, nsp7L8 and the Poly (A) template was incubated 5 min at room temperature to form the active complex. For each assay, 20 μL of this enzyme mix was distributed in wells using a Biomek 4000 (Beckman), containing 2 μL of the compounds. Reactions were initiated by addition of 18 μL of the nucleotide mix (500 μM UTP) and incubated at 30 °C for 20 min. Assays were stopped by the addition of 20 μL of EDTA (100 mM). Reaction mixes were transferred to a Greiner plate using a Biomek I5 Automate (Beckman). Picogreen fluorescent reagent was diluted to 1/800 in TE buffer according to manufacturer’s data, and 60 μL of reagent was distributed into each well of the Greiner plate. The plate was incubated for 5 min in the dark at room temperature, and the fluorescence signal was read at 480 nm (excitation) and 530 nm (emission) using a TecanSafire2.

Positive and negative controls consisted respectively of a reaction mix with 5% DMSO final concentration or EDTA 100 mM or Hinokiflavone 10 µM instead of compounds. For each compound, the percentage of inhibition was calculated as follows: Inhibition % = 100 (raw_data_of_compound − av(pos)/(av(neg) − av(pos)).

Compounds leading to a 30% inhibition or more at 20 µM were selected to further investigations.

The Z’ factor is calculated using the following equation: Z’ = 1 – [3(SD of max) + 3(min SD)] /[(mean max signal) - (mean min signal)], where SD is the standard deviation.

### 2.5 IC_50_ determination based on the fluorescent Picogreen® assay

The compound’s concentration leading to a 50% inhibition of polymerase mediated RNA synthesis was determined in IC_50_ buffer (50 mM HEPES pH 8.0, 10 mM KCl, 2 mM MnCl_2_, 2 mM MgCl_2_, 10 mM DTT) containing 350 nM of Poly(A) template, 150 nM of nsp12 in complex with 1.5 µM nsp7L8 using 7 different concentrations of compound. Five ranges of inhibitor concentration were available (0,01 to 5 µM / 0,1 to 50 µM / 0,5 to 50 µM / 1 to 100 µM / 5 to 400 µM). According to the inhibitory potency of the compound tested, a range was selected to determine the IC_50_. Reactions were conducted in 40 µL on a 96-well Nunc plate. All experiments were robotized by using a BioMek 4000 automate (Beckman). 2 µl of each diluted compound in 100% DMSO were added in wells to the chosen concentration (5% DMSO final concentration). For each assay, the enzyme mix was distributed in wells after a 5 min incubation at room temperature to form the active complex. Reactions were started by the addition of the UTP mix and were incubated at 30°C for 20 min. Reaction assays were stopped by the addition of 20 µl EDTA 100 mM. Positive and negative controls consisted respectively of a reaction mix with 5% DMSO final concentration or EDTA 100 mM instead of compounds. Reaction mixes were then transferred to Greiner plate using a Biomek I5 automate (Beckman). Picogreen***®*** fluorescent reagent was diluted to 1/800° in TE buffer according to the data manufacturer and 60 µl of reagent were distributed into each well of the Greiner plate. The plate was incubated 5 min in the dark at room temperature and the fluorescence signal was then read at 480 nm (excitation) and 530 nm (emission) using a TecanSafire2.

IC_50_ was determined using the equation: % of active enzyme = 100/(1+(I)^2^/IC_50_), where I is the concentration of inhibitor and 100% of activity is the fluorescence intensity without inhibitor. IC_50_ was determined from curve-fitting using Prism software. For each value, results were obtained using triplicate in a single experiment.

## 3. Results

### 3.1 Determination of the experimental conditions of the SARS-CoV RTC fluorescent assay

The nsp12 RdRp activity is dependent on the presence of the nsp7 and nsp8 proteins (Subissi et al., 2014). We made use of a 1:10 ratio of nsp12:nsp7L8 which provides appropriate levels of RdRp activity. A HTS assay was developed with a non-radioactive readout. We used conditions similar to the HTS Dengue Virus (DV) polymerase assay previously developed on our platform. Briefly, the homo-polymeric NTP template was incubated with nsp12:nsp7L8, and the polymerase activity is detected through a fluorescent dye which intercalates upon the synthesis of double stranded RNA (dsRNA). Nsp12 protein and nsp7L8 complex were expressed and purified separately (Fig S1).

We first evaluated the effect of the poly(A) concentration. Maximal SARS-CoV RTC activity was obtained with a Poly(A) concentration of 350 nM, allowing a stable and reproducible signal (Fig. 1A). To define the optimal UTP concentration, the velocity of the RTC was assessed at different UTP concentrations. We observed a non Michaelis-Menten curve with a significant lag-phase (Fig. 1B, full line), and therefore it was not possible to caculate a reliable Km concentration for the robotized assay using standard kinetic modeling. Using a Hill equation fit, we observed an allosteric cooperation with a Hill coefficient of 1.72 (Fig. 1B, dotted line). The UTP concentration to be used was defined at 500 µM, as it corresponds to the beginning of the saturation phase. Finally, to reach 80% of the maximum activity, the nsp12 concentration was set to 150 nM (Fig. 1C).

**Fig. 1.**
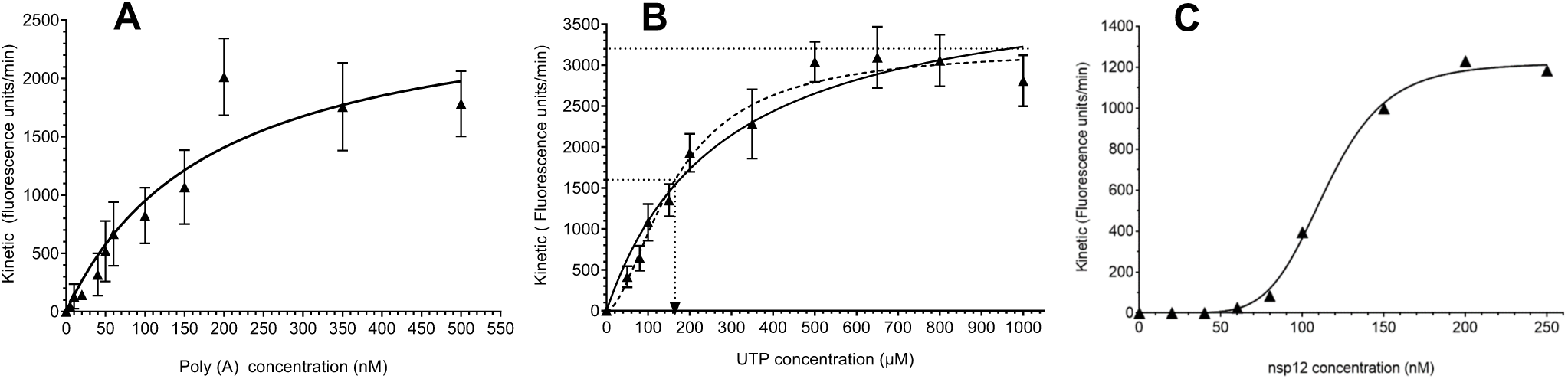
Setting up of the SARS-CoV-RTC activity experimental conditions based on a fluorescent readout. **(A)** Poly(A) template variation with SARS nsp12 in complex with nsp7L8 Velocity values of SARS nsp12 (150 nM) in complex with nsp7L8 (1.5 µM) was measured for various concentrations in Poly (A) template (5; 10; 20; 40; 60; 80; 100; 150; 200; 350; 500 nM). Using the Prism software, velocity values of each condition were determined by calculating the slope of the linear phase of the kinetic and then plotted against the Poly(A) template concentration by using Michaelis-Menten fitting. Data were the results of three independent experiments. **(B)** Nucleotide variation with SARS nsp12 in complex with nsp7L8 Velocity values of SARS nsp12 (150 nM) in complex with nsp7L8 (1.5 µM) was measured for various concentrations in UTP (50;80;100;150;200;350;500;650;800 and 1000 nM). Using the Prism software, velocity values of each condition were determined by calculating the slope of the linear phase of the kinetic and then plotted against the UTP concentration by using Michaelis-Menten equation (full line) or Hill equation (dot line) to determine the apparent Km (UTP) of the SARS-CoV-RTC. Data were the results of three independent experiments. **(C)** Variation in SARS nsp12 in complex with nsp7L8 Velocity values obtained during a time course with 350 nM Poly (A) and 500 µM UTP were plotted against different concentrations in SARS nsp12 in complex with nsp7L8. The concentration ratio nsp12:nsp7L8 was conserved at 1:10. Curve was fitted according the Hill equation.

Using the conditions established above (350 nM poly(A) template, 500 µM UTP with 150 nM nsp12 and 1.5 µM nsp7L8) a time course up to 60 min was performed (Fig. 2). In contrast to the DV2 polymerase, the SARS-CoV RTC complex exhibits a significant lag phase (Fig. 2). To obtain sufficient levels of activity, a reaction time of 20 min. was determined to be suitable for inhibitor screening. As expected, nsp12 alone, or nsp7L8 alone, in the same conditions, did not exhibit any polymerase activity (Fig. 2).

**Fig. 2.**
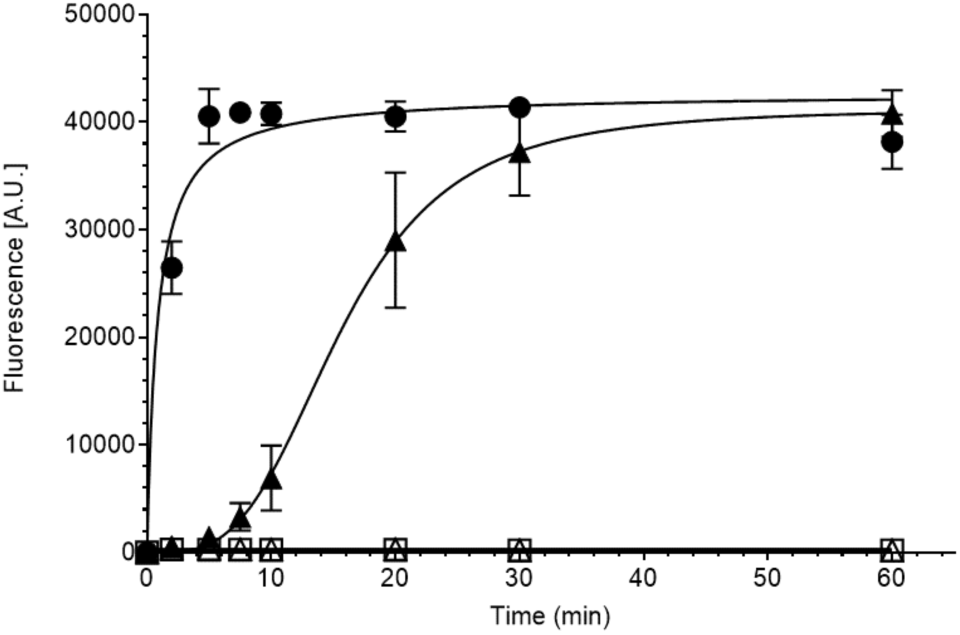
SARS nsp12 in complex with nsp7L8 polymerase activity on a Picogreen kinetic assay. The polymerase activity of 100 nM DV2-NS5 pol (○), 150 nM ns12 in complex with 1.5 µM nsp7L8 (▴), 150 nM nsp12 alone (Δ) or 1.5 µM nsp7L8 alone (□) were measured in a time course (0;2;5;7.5;10;20;30 and 60 min). The produced doubled strand RNA was detected by adding an intercalant reagent (Picogreen®) and by measuring the fluorescence emission at 530 nM. Each assay was performed three times (mean value ± SD).

### 3.2 IC_50_ determination of reference inhibitors of the SARS-CoV RTC

To develop the HTS of the SARS-CoV RTC, reference inhibitors had to be selected. In absence of specific inhibitors of the SARS-CoV RTC we first tested a nucleotide analogue (NA), 3’dUTP, along with a large spectrum of non-specific RNA synthesis inhibitors: Hinokiflavone, Amentoflavone, Quercetin and Apigenin (Coulerie et al., 2012). IC_50_s values range from 0.95 µM for Hinokiflavone to 96 µM for Apigenin (Fig. 3), demonstrating the discriminating capacity of the HTS assay. The 3’dUTP IC_50_ is measured at 6.7 µM under our experimental conditions.

**Fig. 3.**
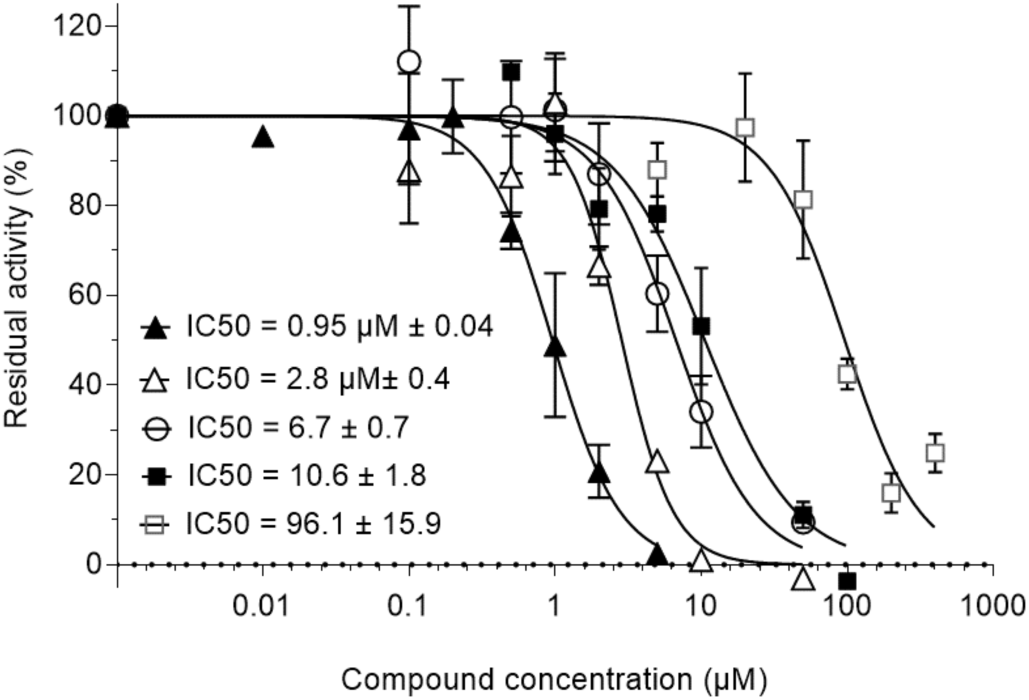
Quantification of SARS-CoV-RTC enzymatic activity inhibitors. Increasing concentrations of Hinokiflavone (▴), Amentoflavone (Δ), 3’dUTP (○), Quercetin (▄) and Apigenin (□) were incubated with 150 nM nsp12, 1.5µ M nsp7Lnsp8, 500 µM UTP, 350 nM Poly (A) at 30°C during 20 min. The Inhibitory concentrations 50 (IC_50_s) were then calculated using graphPad Prism equation (Experiments were done twice in triplicate; Mean value±SD).

### 3.3 High-throughput screening

The total of 1,520 compounds from the Prestwick Chemical Library® (PCL) were tested using the SARS-CoV-RTC assay. The Prestwick Chemical Library **®** is a unique collection of small molecules, mostly approved drugs (FDA, EMA and other agencies) selected by a team of medicinal chemists and pharmacists for their high chemical and pharmacological diversity as well as for their known bioavailability and safety in humans.

The Z’ value was calculated based on the ten control wells on each microplate, resulting in an overall Z’-score of 0,8 +/-0,06 for all the chemical library used in the screen. The screening was performed using 20 µM of compounds in 5% DMSO in a single assay. Based on a cut-off of 30% inhibition, the calculated hit rate is 3.02% (46 hits) and the repartition of identified compounds according their percentage of inhibition was indicated in Figure 4. Based on these results, three main families of molecules were identified: anthracyclines, tetracyclines and detergents, the two first of which were further investigated (Tables A1 and A2).

**Fig. 4.**
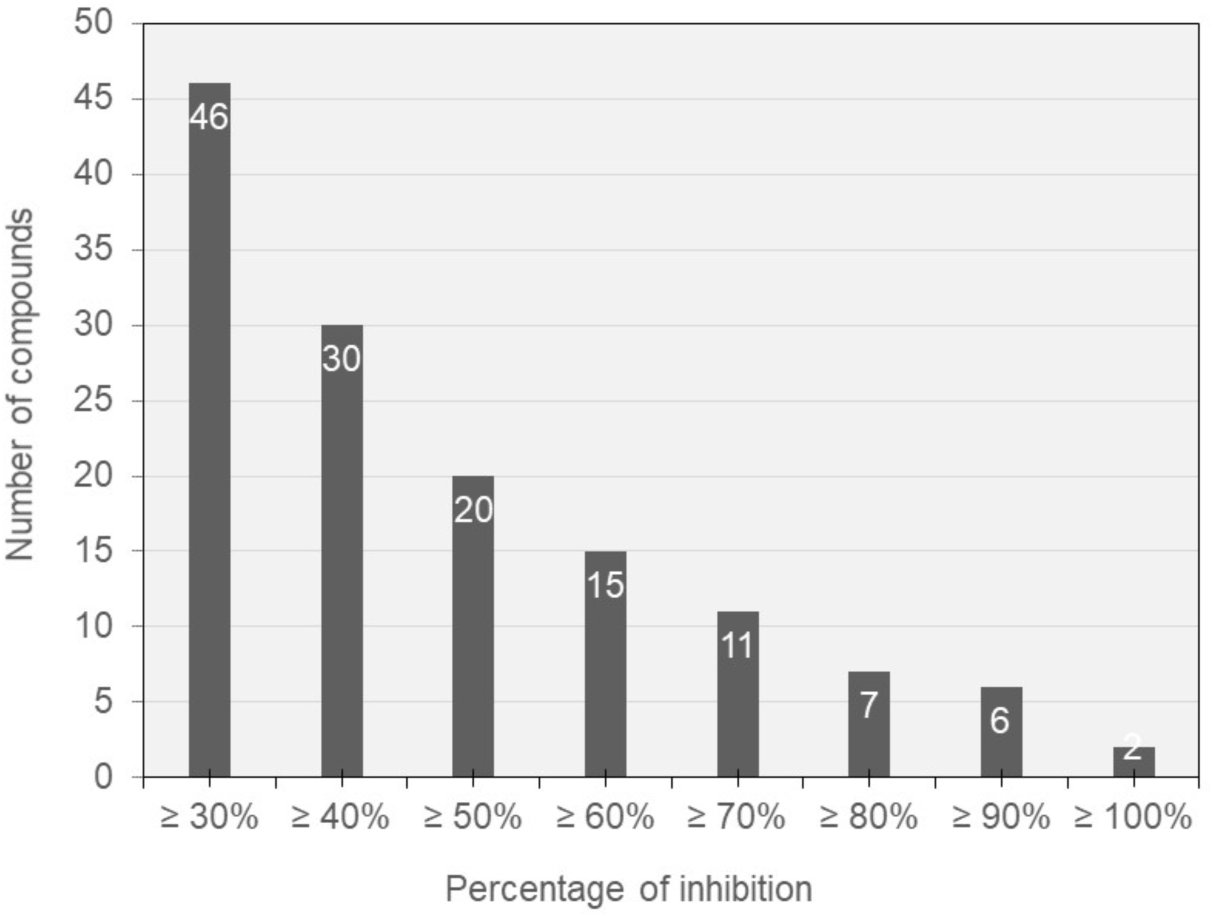
Number of compounds from PCL on the screening of SARS-CoV RTC, according to their inhibitory potential. Based on their efficiency on the SARS-CoV RTC assay, the number of compounds with more of 30%; 40%; 50%; 60%; 70%; 80%; 90% and 100% inhibition were evaluated. The exact value was indicated in white above each bar graph. (Only compounds with more 30% inhibition of the polymerase activity in the assay were represented. Frequent Hitter and fluorescent compounds were excluded). The Z’ value was calculated based on the ten control wells on each microplate, resulting in an overall Z’ score of 0,8 ± 0,06.

### 3.4 IC_50_ determination

Eight anthracyclins were identified with inhibitory potential from zero to 100% at 20 µM (S table 1). Regarding the five determined IC_50_s, they perfectly correlated with the calculated percentage of inhibition at 20 µM of the HTS. Four of the five anthracyclins tested were in the same range of IC_50_s (from 15 to 44 µM). The remaining one, Prestw-385, exhibited a sub-micromolar IC_50_ (0.34 µM ± 0.05) which was in the same range than Hinokiflavone, the inhibitor control (Fig. 5A).

**Fig. 5.**
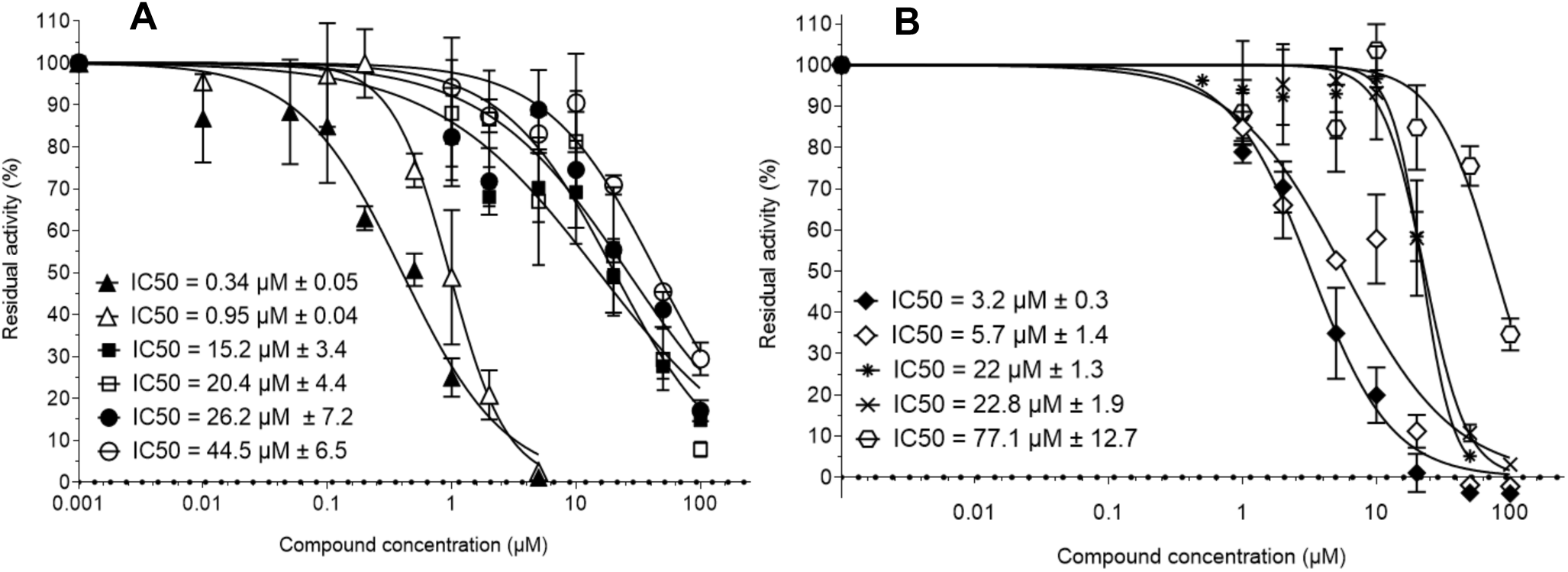
Quantification of SARS-CoV-RTC enzymatic activity inhibitors belonging to the anthracycline and tetracyclin chemical families. **(A)** Anthracyclin chemical family. Increasing concentrations of Prestw-385 (▴), Prestw-1752 (▄), Prestw-1224 (□), Prestw-438 (○) and Prestw-487 (○) were used to determine the IC_50_s of each compound. Hinokiflavone (Δ) was indicated as a control. **(B)** Tetracycline chemical family. Increasing concentrations of Prestw-456 (♦), Prestw-1799 (◊), Prestw-1000 (*), Prestw-145 (X) and Prestw-140 () were used to determine the IC_50_s of each compound. Compounds were incubated with 150 nM nsp12, 1.5 µM nsp7Lnsp8, 500 µM UTP, 350 nM Poly (A) at 30°C during 20 min. The IC_50s_ were then calculated using graphPad Prism equation (Experiment were done twice in triplicate; Mean value ± SD). IC_50_: concentration for 50% inhibition. The IC_50_ of Prestw-140 was approximate.

Ten tetracyclins were identified in the PCL, with inhibitory potential up to 74% at 20 µM (S table 2). Again, the calculated IC50 values for these compounds was found to perfectly correlate to the percentage of inhibition at 20 µM. Three tetracyclins showed IC_50_s of 20 µM or higher (Prestw-1000; Prestw-145; Prestw-140) while Prestw-456 and Prestw-1799 exhibited lower IC_50_s of 3.2 and 5.7 µM respectively (Figure 5B).

## 4. Discussion

In this study we have established a robust HTS assay with a fluorescent readout for the SARS-CoV RTC. The overall hit rate for the PCL screen is ∼3%, with a cut-off of 30% of inhibition. Obviously, at this stage the identified hits are active only on the SARS-CoV RTC and have by no means been deemed suitable for medication.

Regarding the selected two families of molecules, the anthracyclins and the tetracyclins, the number of identified molecules did not allow Structure Activity Relationship (SAR) studies. Rather, it is important to outline the accuracy of the IC_50_s of the primary screening, as well as the reproducibility of the IC50s regarding the calculated standard errors. With the exception of the anthracyclins, which have been described as potential intercalating molecules, it is not known at this stage if these compounds are indeed intercalating, denaturing, or low specificity agents having a mechanism of action suitable for advanced drug design. Preliminary testing in SARS-CoV-2 infected cells indicate that none of the compounds described here show significant antiviral activity (data not shown).

In summary, this work, to the best of our knowledge, is the first description of a robust HTS assay based on a SARS-CoV RTC. It provides a new strategy for the rapid identification of potential anti-SARS inhibitors. The next steps already under development include increasing the output by scaling up the assay to 384 and potentially 1536 wells plates. Furthermore, the screening of large chemical libraries derived from innovative approaches as virtual screenings or Protein Protein Interaction inhibitors (iPPI) will broaden the scope of potential antivirals and constitute a key step to speed up these objectives. Identification of candidate anti-coronavirus drugs, based on a diversity of chemical libraries, should expedite drug discovery and design, and constitute an invaluable help to confirm the target of hits identified during cell-based or phenotypic screens.

## Acknowledgements

This work was supported by the Fondation pour la Recherche Médicale (Aide aux équipes), the SCORE project H2020 SC1-PHE-Coronavirus-2020 (grant#101003627), Inserm through the REACTing initiative (REsearch and ACTion targeting emerging infectious diseases), and the ANR-Flash-COVID (ANR-20-COVI-0047-01, TAMAC), supported by the Fondation de France. We thank ML Jung, E. Decroly for helpful comments.

---

**Fig. A1.**
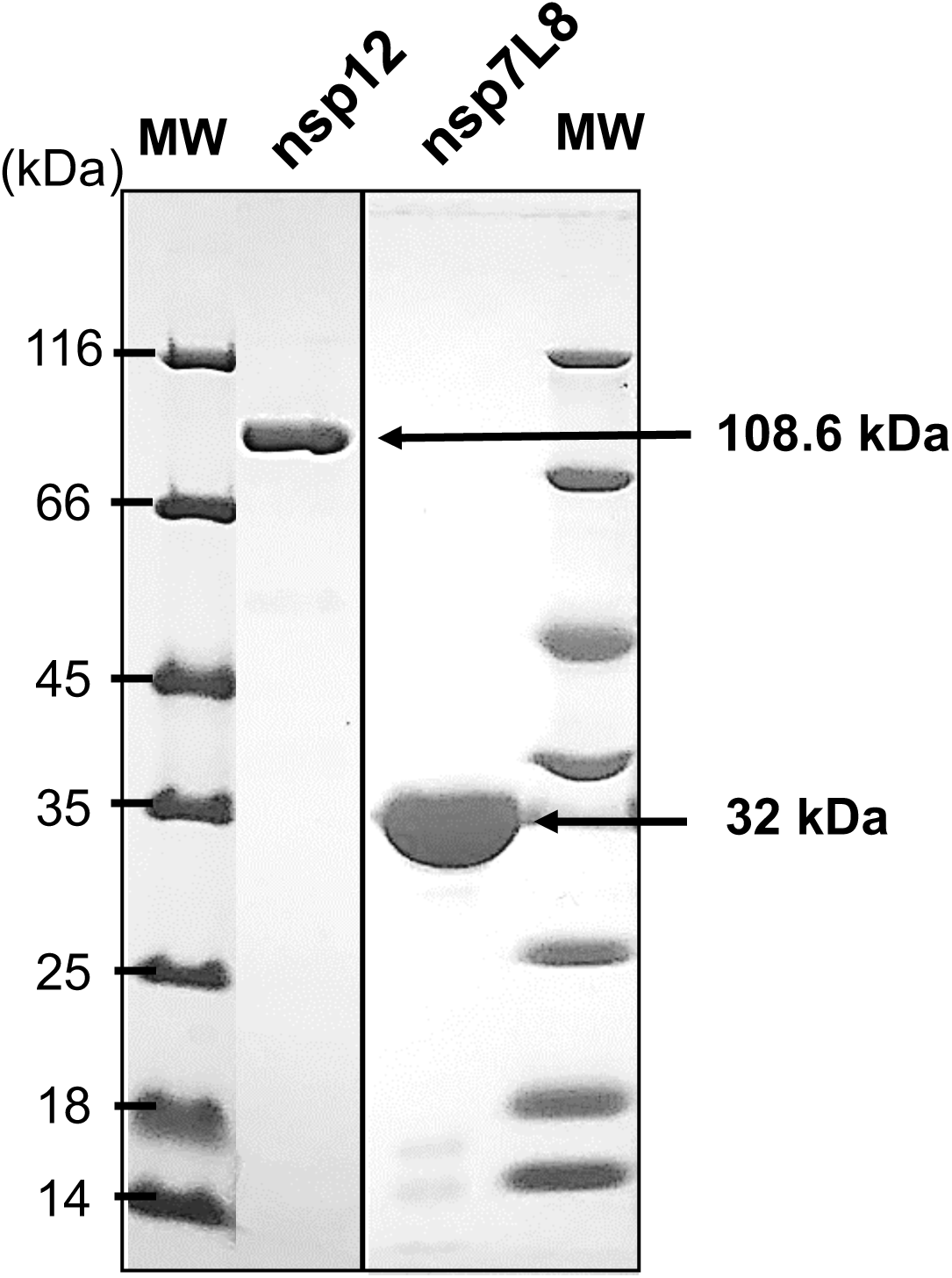
Characterization of the purified recombinant SARS nsp12 and nsp7L8 proteins. 5 µg of SARS nsp12 and 5 µg of nsp7L8 were loaded on a 10% and 12% SDS-PAGE gel respectively, stained by Coomassie blue. (MW, molecular size markers)

**Table A1.** Percentage of inhibition and Inhibitory Concentration 50 of compounds related to the anthracyclin chemical family. The compounds are listed according to their inhibitory effect (N.D.: Not Determined).

| Compound | % Inhibition at 20 μM | IC<sub>50</sub> (μM) |
| --- | --- | --- |
| Prestw-385 | 101.12% | 0.34 ± 0.06 |
| Prestw-1752 | 41.13% | 15.2 ± 3.4 |
| Prestw-1224 | 50.08% | 20.4 ± 4.4 |
| Prestw-438 | 52.12% | 26.2 ± 7.2 |
| Prestw-487 | 64.84% | 44.5 ± 6.5 |
| Prestw-1399 | 23.72% | N.D. |
| Prestw-1167 | 17.63% | N.D. |
| Prestw-1703 | -20.5% | N.D. |

**Table A.2.**
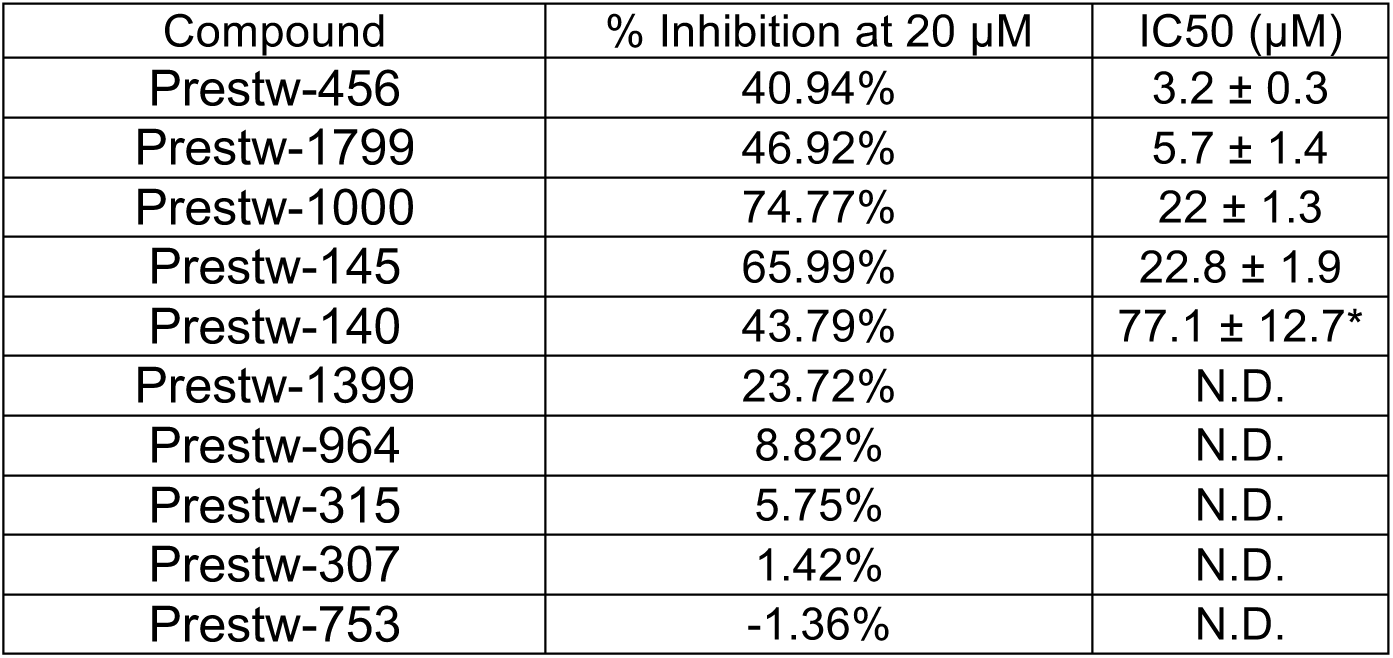
Percentage of inhibition and Inhibitory Concentration 50 of compounds related to the tetracyclin chemical family. The compounds are listed according to their inhibitory effect. (* approximate IC_50_): (N.D.: Not Determined).

| Compound | % Inhibition at 20 μM | IC<sub>50</sub> (μM) |
| --- | --- | --- |
| Prestw-456 | 40.94% | 3.2 ± 0.3 |
| Prestw-1799 | 46.92% | 5.7 ± 1.4 |
| Prestw-1000 | 74.77% | 22 ± 1.3 |
| Prestw-145 | 65.99% | 22.8 ± 1.9 |
| Prestw-140 | 43.79% | 77.1 ± 12.7* |
| Prestw-1399 | 23.72% | N.D. |
| Prestw-964 | 8.82% | N.D. |
| Prestw-315 | 5.75% | N.D. |
| Prestw-307 | 1.42% | N.D. |
| Prestw-753 | -1.36% | N.D. |

